# Complete guide RNA design for CRISPR-mediated regulation of human long noncoding RNA transcription

**DOI:** 10.1101/290338

**Authors:** Amir Saberi, Renjun Zhu, Chulan Kwon

**Affiliations:** Division of Cardiology, Johns Hopkins University School of Medicine

## Abstract

Transcription inhibition and activation of long noncoding RNAs (lncRNAs) mediated by clustered regularly interspaced short palindromic repeat (CRISPR)/Cas9 technology provides potential advantages in high-throughput functional genomics studies over RNA interference or overexpression platforms. In this work, we identify over 90,000 lncRNA transcription start sites (TSSs) based on the MiTranscriptome human genome annotation and design single guide RNA (sgRNA) libraries with strong predicted activities and low off-target effects for CRISPR-mediated inhibition and activation (CRISPRi/a) of their transcription. A large fraction of these TSSs correspond to putative genes that are not annotated in common reference genome annotations and have never been functionally studied. Our CRISPRi/a libraries, or their context-dependent subsets, are potentially useful in genome-scale functional studies of human lncRNAs.

## Introduction

Genetic programming platforms capable of precise regulation of gene transcription are essential for development of novel cell engineering technologies and rigorous interrogation of complex biological systems. Current methods of gene regulation have intrinsic limitations that render them unsuitable for scaling and complicate interpretation of their outcomes. For example, gene silencing by RNA interference (RNAi) can have large off-target effects and unpredictable efficiencies that depend on subcellular localization of the target transcript, among other gene-specific factors (Fedorov et al. 2006; Mohr et al. 2014). Protein-based tools such as transcription-activator-like effectors (TALEs) fused to activators or repressors allow site-specific regulation of transcription, but are difficult to design, build, and deliver. Recently, catalytically inactive mutants of the CRISPR-related Cas9 enzyme (dCas9) have been used to activate or repress gene expression directly from genomic loci by RNA-guided targeting of transcription factors and chromatin modifiers. This method, known as CRISPR interference/activation (CRISPRi/a), provides an efficient, robust, multiplexable, and programmable approach for genome-wide regulation of expression (Gilbert et al. 2013; Gilbert et al. 2014; Qi et al. 2013; Tanenbaum et al. 2014; Zalatan et al. 2015).

CRISPRi/a is especially advantageous for regulation of non-coding RNA expression since these genes lack an open reading frame (ORF) and their function is relatively unaffected by small CRISPR-induced insertion/deletion modifications. Furthermore, a large fraction of them reside in the nucleus, rendering them immune to RNAi (Goyal et al. 2017). Long non-coding RNAs (lncRNAs) are emerging as potent regulators of cellular function (Ponting et al. 2009; Fatica & Bozzoni 2014; Wang & Chang 2011). However, only a small fraction of annotated lncRNAs have been functionally studied and a vast majority of transcribed lncRNAs remain unannotated, and thus, excluded from genome-wide and other studies. While the most recent human genome annotation includes fewer than 16,000 lncRNAs (Harrow et al. 2012), a meta-analysis of human healthy and diseased tissue and cell line transcriptomes has identified close to 60,000 lncRNA genes that comprise over 68% of human genes (Iyer et al. 2015). Characterization of this enormous collection of virtually unknown genes is essential to understanding normal and pathological human biology.

In this work, we aimed to identify single guide RNA (sgRNA) target sites that enable transcription activation or repression of nearly every lncRNA locus in the human genome. Using a previously developed machine learning approach optimized using experimental data (Horlbeck et al., 2016), we identified sets of highly active sgRNAs with minimal predicted off-target effects for activation or repression of over 90 thousand human lncRNAs. These sgRNA libraries or context-dependent subsets of them can be used in high-through screening platforms to uncover the biological function of previously unknown or understudied human lncRNAs.

## Results

To design sgRNAs for transcriptional regulation of human lncRNAs, we sought to identify nearly all lncRNA transcription start sites (TSSs) across the genome. To this end, we turned to MiTranscriptome version 2.0, a catalog of over 91 thousand human long polyadenylated RNAs that are mostly uncharacterized and unannotated in RefSeq, UCSC, or GENCODE (Iyer et al. 2015). Of these, 58,648 genes correspond to lncRNAs that are transcribed from 99,760 unique TSSs. About 11% of these TSSs were in proximity of those identified by cap analysis of gene expression (CAGE) (Kodzius et al. 2006). In these cases, the CAGE-based TSS annotation was used in our sgRNA design, while the rest of the MiTranscriptome TSS annotations were directly used (Figure 1A).

**Figure 1.**
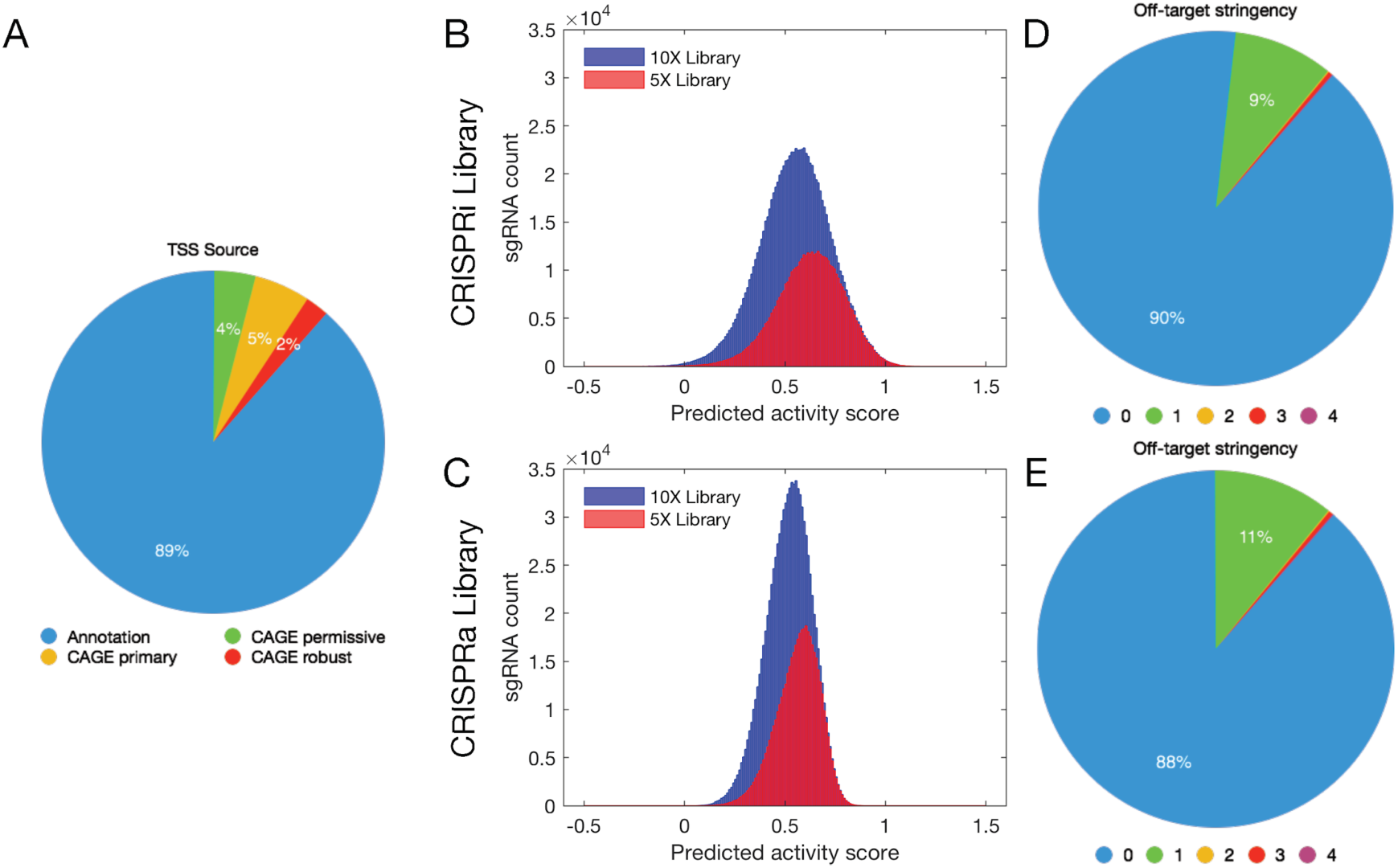
CRISPRi/a libraries for human lncRNAs. (A) Pie chart showing the composition of the source of lncRNA transcription start sites (TSSs). The majority of TSSs were derived from MiTranscriptome annotations, while others were derived by matching to the CAGE-based FANTOM5 expression atlas. (B–C) Histograms of predicted activity scores of CRISPRi (B) and CRISPRa (C) libraries. Blue histograms correspond to 10X libraries, while red histograms show the compact 5X libraries. (D–E) Pie charts showing off-target stringencies for CRISPRi (D) and CRISPRa (E) sgRNA libraries. Higher off-target stringency values indicate more relaxed conditions in off-target filtering.

For sgRNA design, we used an integrated machine learning approach developed previously for prediction of activity in CRISPRi and CRISPRa (Horlbeck et al. 2016). This method integrates several parameters including sgRNA size and composition, distance to TSS, and predicted secondary RNA structure to calculate a score for each potential protospacer sequence regarding its activity in CRISPRi/a. To facilitate the computational task, we divided the TSS set into chunks of maximum 20,000 loci that were then analyzed by the algorithm to predict the best 5 and the next best 5 sgRNAs for CRISPRi and CRISPRa activity. Among all lncRNA TSSs, we were able to create CRISPRi sgRNAs for 97,627 and CRISPRa sgRNAs for 94,644, while for other TSSs there was not sufficient number of potential protospacer sequences passing all design criteria. The sgRNA subsets for interference and activation were then separately pooled to yield the whole-genome CRISPRi/a sgRNA libraries (available at https://www.dropbox.com/sh/snzj68vl99iyx8t/AADO9WDH4mRYl7CpISnGdInda).

The median predicted activity score for our 10X CRISPRi library is 0.557, with the upper and lower quartiles of 0.671 and 0.439, respectively. When including only the top 5 sgRNAs (5X library), the median activity score was improved to 0.633 with the upper and lower quartiles of 0.741 and 0.520 (Figure 1B). Our CRISPRa libraries have slightly lower median predicted activity scores (0.526 and 0.575 for 10X and 5X CRISPRa library, respectively), but slightly tighter distribution (upper and lower quartiles 0.600 and 0.446 for the 10X library and 0.641 and 0.497 for the 5X library, Figure 1C). Of all sgRNAs, only a small percentage (19% and 15% for 10X CRISPRi/a libraries, respectively) have a predicted scores less than 0.4, which indicate inactive sgRNAs. These numbers further reduced to 9% and 7% in the 5X CRISPRi/a libraries. On the other hand, a large population of sgRNAs (40% and 25% for 10X CRISPRi/a libraries, respectively) have activity scores greater than 0.6, indicating active sgRNAs. These numbers further improved to 58% and 40% in the 5X libraries. More importantly, sgRNAs with predicted scores greater than 0.6 cover the vast majority of TSSs (89% and 73% for 10X CRISPRi/a libraries, 85% and 70% for 5X CRISPRi/a libraries, respectively), suggesting there is at least 1 sgRNA with strong predicted activity for the majority of TSSs. The slight reduction in highly active sgRNA TSS coverage in the 5X libraries is due to exclusion of some sgRNAs with high activity scores but also high predicted off-target effect. A vast majority of our predicted CRISPRi/a sgRNAs (90% for interference and 88% for activation 10X libraries) have low predicted off-target activities indicated by the low off-target stringency (Figure 1D and E).

## Conclusions

Despite their emerging fundamental roles in regulation of cellular function, human the vast majority of human lncRNAs remain uncharacterized or absent from public annotation databases altogether. Systematic study of lncRNA function in normal and diseased conditions will benefit from development of novel efficient and scalable gene perturbation platforms. RNA-guided CRISPR-mediated gene activation and repression technologies using effector-conjugated dCas9 proteins have the potential to enable such genetic programming platforms. However, their utility depends on identification of potent and specific sgRNAs for the set of target genes. In this work, we identified CRISPRi/a sgRNA target sites with high predicted activity, low predicted off-target effects, and near-complete coverage for the largely uncharacterized set of over 90 thousand human lncRNA TSS loci. These libraries or their context-dependent subsets can potentially enable future genome-scale lncRNA screens or novel cell and tissue engineering approaches.

## References

Fatica, A. & Bozzoni, I., 2014. Long non-coding RNAs: new players in cell differentiation and development. Nature reviews. Genetics, 15(1), pp. 7–21.

Fedorov, Y. et al., 2006. Off-target effects by siRNA can induce toxic phenotype. RNA, 12(7), pp. 1188–1196.

Gilbert, L.A. et al., 2013. CRISPR-mediated modular RNA-guided regulation of transcription in eukaryotes. Cell, 154(2), pp. 442–451.

Gilbert, L.A. et al., 2014. Genome-Scale CRISPR-Mediated Control of Gene Repression and Activation. Cell, 159(3), pp. 647–661.

Goyal, A. et al., 2017. Challenges of CRISPR/Cas9 applications for long non-coding RNA genes. Nucleic acids research, 45(3), p.e12.

Harrow, J. et al., 2012. GENCODE: the reference human genome annotation for The ENCODE Project. Genome research, 22(9), pp. 1760–1774.

Horlbeck, M.A. et al., 2016. Compact and highly active next-generation libraries for CRISPR-mediated gene repression and activation. eLife, 5. Available at: http://dx.doi.org/10.7554/eLife.19760.

Iyer, M.K. et al., 2015. The landscape of long noncoding RNAs in the human transcriptome. Nature genetics, 47(3), pp. 199–208.

Kodzius, R. et al., 2006. CAGE: cap analysis of gene expression. Nature methods, 3(3), pp. 211–222.

Mohr, S.E. et al., 2014. RNAi screening comes of age: improved techniques and complementary approaches. Nature reviews. Molecular cell biology, 15(9), pp. 591–600.

Ponting, C.P., Oliver, P.L. & Reik, W., 2009. Evolution and Functions of Long Noncoding RNAs. Cell, 136(4), pp. 629–641.

Qi, L.S. et al., 2013. Repurposing CRISPR as an RNA-guided platform for sequence-specific control of gene expression. Cell, 152(5), pp. 1173–1183.

Tanenbaum, M.E. et al., 2014. A protein-tagging system for signal amplification in gene expression and fluorescence imaging. Cell, 159(3), pp. 635–646.

Wang, K.C. & Chang, H.Y., 2011. Molecular Mechanisms of Long Noncoding RNAs. Molecular cell, 43(6), pp. 904–914.

Zalatan, J.G. et al., 2015. Engineering complex synthetic transcriptional programs with CRISPR RNA scaffolds. Cell, 160(1-2), pp.339–350.

